# MORPHOGENIC VERSUS MITOGENIC ROLES OF SHH ARE SEGREGATED ON DISTINCT EXOSOMES REGULATED BY CELLULAR RAB7 LEVELS

**DOI:** 10.1101/2023.06.27.546648

**Authors:** Ankita Walvekar, Shivangi Pandey, Siddhesh S. Kamat, Sharat Damodar, Raj K. Ladher, Neha Vyas

**Author notes:** **Corresponding authors:** 1. Dr. Neha Vyas, Assistant Professor, Division of Molecular Medicine, St. John’s Research Institute, St. John’s National Academy of Health Sciences, Bangalore - 560 034., Karnataka, India., 2. Dr. Raj K. Ladher, National Centre for Biological Sciences, GKVK, Bellary Road, Bangalore 560065., Karnataka, India.

## Abstract

The secreted signaling molecule, Sonic hedgehog (Shh) is involved in patterning and growth of various embryonic tissues across species. This lipid-anchored protein is secreted on extracellular vesicles to activate signaling. Shh has two functions during spinal cord development, acting as a morphogen to pattern the ventral neural tube and a mitogen to maintain neural progenitors. Here, we find that these activities are segregated on to two distinct pools of exosomes. Shh secreted on a classical exosomal pool (Shh-P150) is able to activate ventral threshold targets in the neural tube. In contrast, the mitogenic activity, is elicited by a lighter exosomal pool (Shh-P450). We further show that cellular environment plays a major role in regulating the biogenesis of P150 versus P450 by modulating Rab7 levels. We find that active Rab7 is necessary for ventral neural tube patterning through the regulation of Shh-P150 secretion. The cellular mechanisms involved in packaging and secretion of Shh with different partners on P150 or P450 pools may be more general, enabling the separation of functional activities of signaling molecules during development and disease.

## Introduction

The Hedgehog (Hh) proteins are a highly conserved family of morphogens. First identified in Drosophila, Hh and its vertebrate counterparts Sonic (Shh), Indian (Ihh), and Desert hedgehog (Dhh) show diverse roles in embryonic tissue patterning (Ingham and McMahon, 2001), tissue-specific stem cell maintenance (Briscoe and Thérond, 2013), tissue homeostasis (Ingham, Nakano and Seger, 2011) and even cancer progression (Taipale and Beachy, 2001; Barakat, Humke and Scott, 2010). Hh is post-translationally lipidated, with cholesterol (Porter *et al*., 1995) and palmitate (Pepinsky *et al*., 1998) moieties added to the N and C-terminal of the processed protein, respectively. These modifications are essential for the precision of Hh signaling (Manikowski, Kastl and Grobe, 2018). Despite being modified by two hydrophobic membrane-anchors, Hh acts in a paracrine manner, activating both short and long-range targets in the developing embryos (Briscoe and Thérond, 2013). There is evidence to suggest that these activities may result from the secretion of Hh through exosomes (Gradilla *et al*., 2014; Matusek *et al*., 2014; Vyas *et al*., 2014; Parchure *et al*., 2015) or on lipoproteins (Panáková *et al*., 2005; Palm *et al*., 2013). For exosomal secretion, the membrane-anchored Hh proteins are trafficked into a specific late-endosomal compartment the multi-vesicular body (MVB), and sorted on intra-luminal vesicles (ILVs). Upon fusion of the MVB to the plasma membrane, Hh containing ILVs are secreted as exosomes to activate the signaling (Raposo and Stoorvogel, 2013; Parchure, Vyas and Mayor, 2017).

Sonic hedgehog (Shh) emanating from the notochord mediates patterning of the ventral regions of the neural tube (Roelink *et al*., 1995; Ericson *et al*., 1996). Here, Shh acts as a classical morphogen by spatially restricting the expression of different transcription factors in subsets of neuronal precursors (Ribes and Briscoe, 2009). It does this by both activating ventral determinants and repressing dorsal ones, in a concentration-dependent fashion (Briscoe *et al*., 2000). Concomitantly, Shh also regulates growth and proliferation in the neural tube. Inhibition of Shh signaling abrogates the proliferation and survival of neural progenitors (Cayuso *et al*., 2006). Conversely, the over activation of Shh signaling pathway results in the over-proliferation of progenitors (Cayuso *et al*., 2006; Ulloa and Briscoe, 2007). Both these roles ensure the precise development of the neural tube, however, the mechanism through which these specific signaling roles of Shh are segregated are unclear.

Our previous work identified two different types of exosomal pools, termed Shh-P150 and Shh-P450, based on their pelletability by ultracentrifugation (Vyas *et al*., 2014). Our work had shown an activity for Shh-P150 exosomes in the differentiation of ventral neuronal progenitors from mouse embryonic stem cells (ES cells), however Shh-P450 appeared to not show an activity in this assay (Vyas *et al*., 2014). Here we further explore the function of these two pools of exosomes in the spinal cord patterning. Using neural explants from early chick embryos, we find that Shh segregates its two activities on two distinct pools of exosomes during neural tube development. Shh carried on P150 exosomes induces the expression of ventral transcription factors. In contrast, while Shh-P450 can induce the transcription of Nkx6.1, a broad early ventral precursor gene, protein expression is not detectable. Instead, Shh-P450 shows a mitogenic activity on neural plate explants. Our findings suggest an intricate cellular process that allows a signaling molecule to toggle between two different activities, in this case morphogenic and mitogenic role, by changing its partners or packaging.

## Results

### Shh-P150 functions as a morphogen during spinal cord development

As previously described, Shh-P150 and Shh-P450 pools were isolated from HEK293T cells expressing Shh (Shh-HEK), by sequential ultracentrifugation^9^. The pellets were resuspended and the concentration of Shh in each fraction was measured using densiometric analysis for each batch (Fig. S1). P150 and P450 pools were also collected from untransfected HEK293T cells, and used as controls for all assays.

In order to determine the activity of the two Shh exosomal pools, we used neural plate explants from the caudal region of HH10 stage chicken embryos. These were embedded in a collagen matrix (Yamada *et al*., 1993) and treated with 4 nM Shh-P150 or 4 nM Shh-P450 for 48 hrs, after initial standardizations (Figs. S2, S3). In these conditions, we found that Shh-P150 was able to induce FoxA2, Nkx2.2 and Nkx6.1 protein expression (Fig. 1). In contrast, Shh-P450 was unable to induce the expression of these ventral neural markers (Fig. 1). Shh signaling has also been shown to repress dorsal determinants of the neural tube. We thus asked if Shh-P150 or Shh-P450 were able to repress Pax7 in the neural explant assay. We found that neither Shh exosomal pool repressed Pax7 expression, although Purmorphamine (a Smoothened agonist) showed robust Pax7 repression (Fig. S4).

**Fig. 1:**
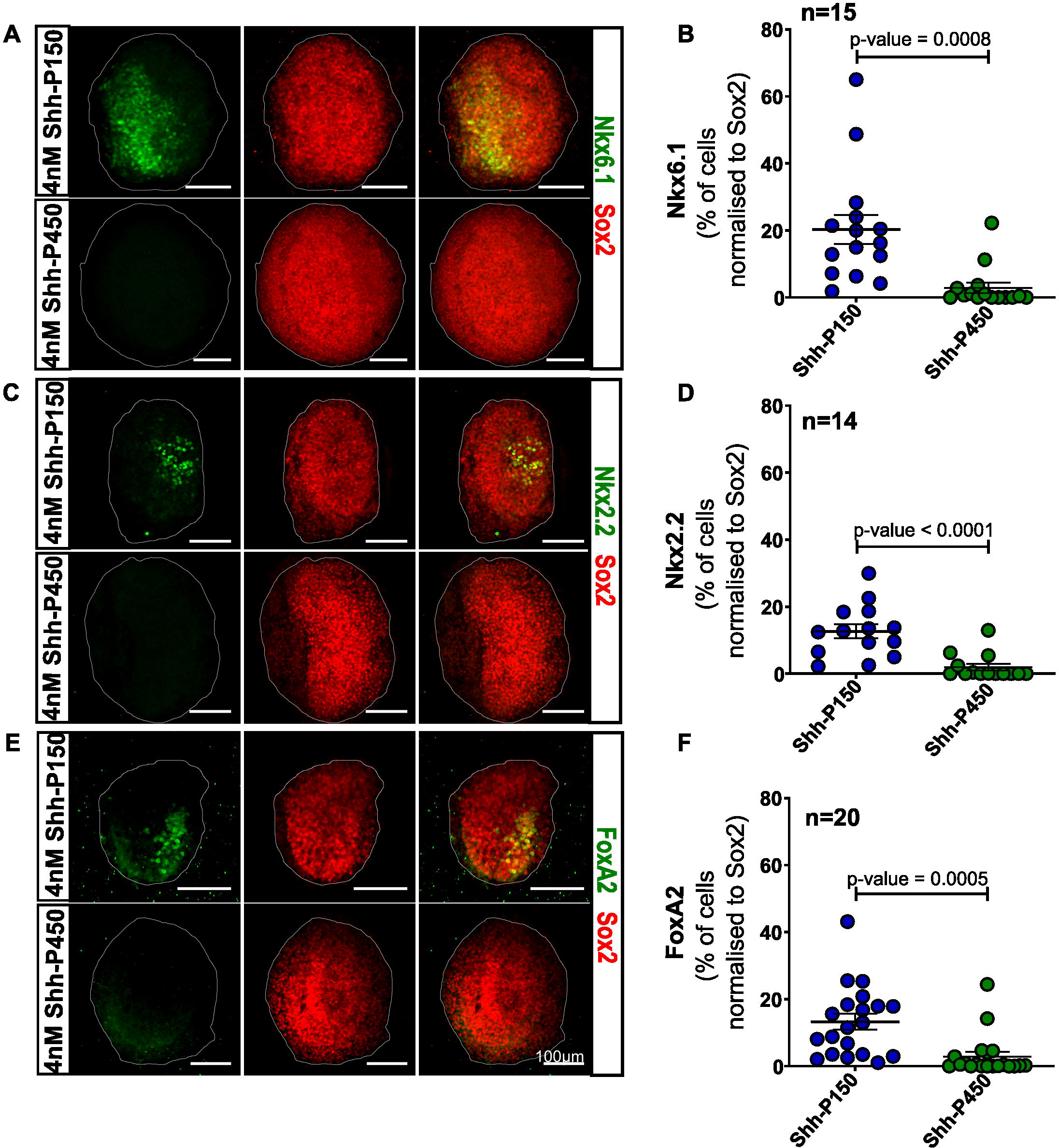
Shh-P450 is inefficient at ventral neural progenitor specification, unlike Shh-P150. (A, C, E) Representative immunofluorescence images of neural plate explant treated with 4 nM Shh-P150 (top panel in A, C, E) and 4 nM Shh-P450 (bottom panel in A, C, E) exosomes for 48 hrs. (A) Nkx6.1 (green, left panel and merge), (C) Nkx2.2 (green, left panel and merge), (E) FoxA2 (green, left panel and merge). Pan neural progenitor marker, Sox2 (red, in all middle panels and merge), scale bar, 100 μm. (B, D, F) Dot plots represent quantitative analysis of the immunofluorescence images of explants incubated with 4 nM Shh-P150 (in blue) and 4 nM Shh-P450 (in green) for 48 hrs. Each data-point represents percentage positive nuclei normalized to Sox2 positive nuclei for each explant, Nkx6.1 (B), Nkx2.2 (D), FoxA2 (F). Error bars represent mean ± SEM. Student’s test was performed to derive p-values. n, number of explants.

To ask if Shh-P450 required a higher concentration to show activity, we incubated explants in 8 nM Shh-P150 or Shh-P450 for 48 hours (Fig. 2A-F). Shh-P150 could induce the expression of FoxA2, Nkx2.2 and Nkx6.1 (Fig. 2A-F, upper panel). However, even at 8 nM concentration Shh-P450 was unable to activate these ventral markers (Fig. 2A-F, lower panel). P150 and P450 pools from untransfected HEK293T cells used as controls did not activate ventral markers at equivalent total protein concentrations of respective Shh exosomal pools for 4 nM or 8 nM (Figs. S5 & S6)

**Fig. 2:**
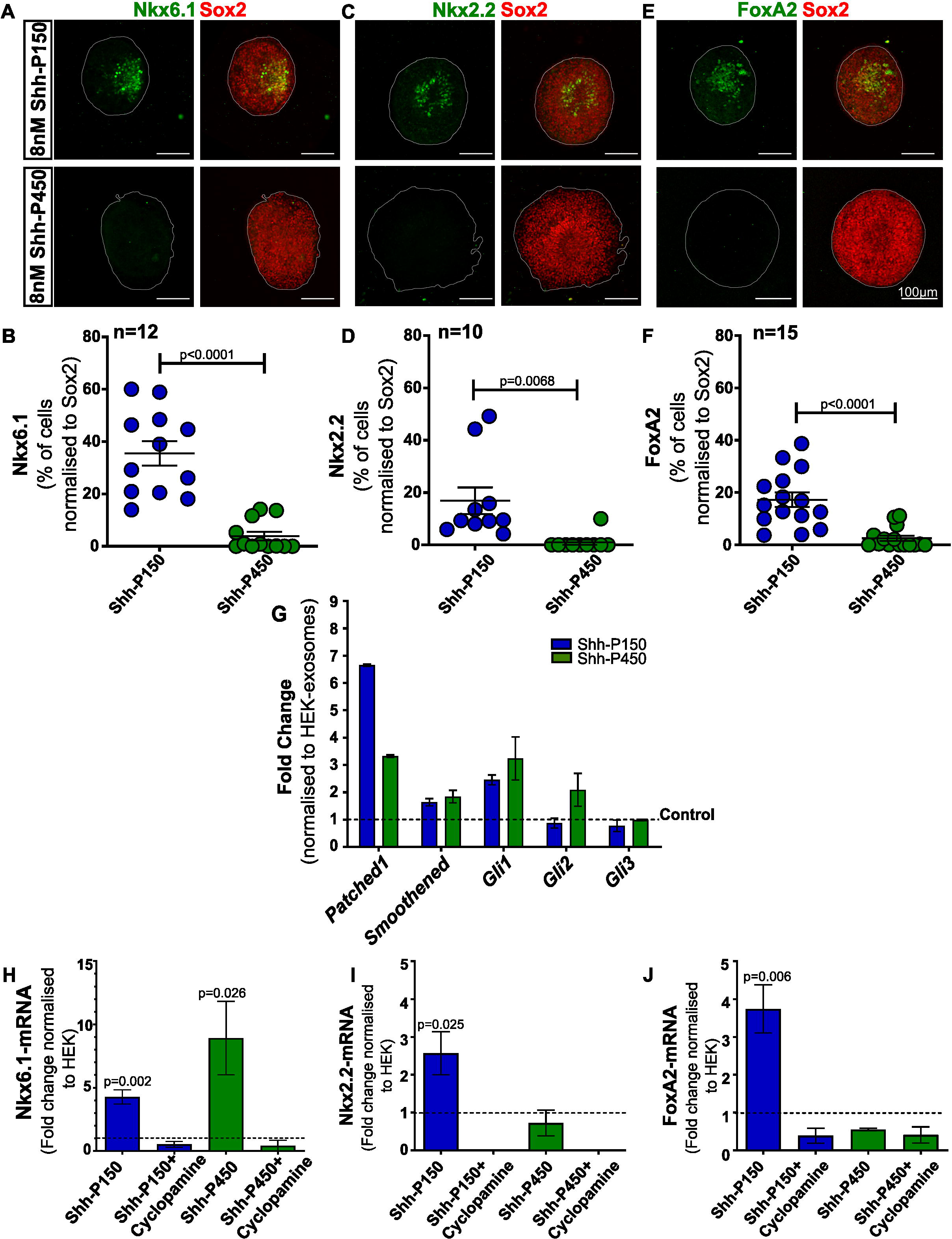
Shh-P450 pool fails to specify ventral neural progenitors but can activate Shh-pathway. (A, C, E) Representative immunofluorescence image of neural explants treated with 8 nM Shh-P150 (top panel) and 8 nM Shh-P450 (bottom panel) for 48 hrs and stained for Nkx6.1 (A, green, left panel and merge), Nkx2.2 (C, green, left panel and merge), and FoxA2 (E, green, left panel and merge). Pan neural progenitor marker, Sox2 (red, all merge). (B, D, F) Dot plots representing percentage of Nkx6.1 (B), Nkx2.2 (D), FoxA2 (F) positive nuclei, normalized to Sox2 positive nuclei. Each dot represents data from one explant. 8 nM Shh-P150 treated explants have higher number of cells with Nkx6.1, Nkx2.2, and FoxA2 expression (blue dots in B, D, F) than 8 nM Shh-P450 treated explants (green dots in B, D, F). Error bars represent mean ± SEM. Student’s test was performed to derive p-values. n, number of explants. (G) qPCR analysis of Shh pathway genes (*Patched1, Smoothened, Gli1, Gli2, Gli3)*, using explants treated with 4 nM Shh-P150 (blue bars) or 4 nM Shh-P450 (green bars). Fold change was derived using explants treated with equal total protein containing HEK-P150 or HEK-P450 pools, respectively. Dotted line represents levels of respective HEK normalization. n = 2 biological replicates. Error bars represent mean ± SEM. (H-J) qPCR analysis of explants treated with 4 nM Shh-P150 (blue bars) or 4 nM Shh-P450 (green bars), with or without 1 μΜ Cyclopamine. Fold change was derived using explants treated with equal total protein containing HEK-P150 or HEK-P450 pools, respectively, represented as dotted line. *Nkx6.1* (H), *Nkx2.2* (I) and *FoxA2* (J). n = 3 biological replicates. Statistical significance determined using Student’s T-test. Error bars represent mean ± SEM.

To examine if the failure of Shh-P450 to induce ventral gene expression was as a result of its inability to trigger the Shh signaling, we evaluated the ability of Shh-P450 pool to activate the Shh-pathway. At 48 hrs, both 4 nM Shh-P150 and 4 nM Shh-P450 pools could activate transcription of endogenous Hh-pathway genes (Fig. 2G). We found that the Shh-P450 pool not only upregulates the transcriptional effector, Gli1 but also Gli2 expression. In contrast, Shh-P150 showed only Gli1 upregulation.

The inability of Shh-P450 to induce ventral marker genes, even though it was able to signal, led us to ask if Shh-P450 could only stimulate weak transcription of ventral neural tube determinants, that were below the detection range of immunohistochemistry. We therefore used qPCR to determine transcript levels of ventral determinants. Consistent with our results from antibody staining, Shh-P150 was able to stimulate *FoxA2*, *Nkx2.2* and *Nkx6.1* expression (Fig. 2H-J, blue bars). While Shh-P450 was unable to stimulate *FoxA2* and *Nkx2.2* transcription (Fig. 2I-J, green bars), *Nkx6.1* transcripts were detected at comparable, if not higher, levels to Shh-P450 (Fig. 2H, green bars). This stimulation was sensitive to cyclopamine (Fig. 2H-J). Taken together, these data show that both Shh-P150 and Shh-P450 pools can signal, however only Shh-P150 is able to induce ventral determinants in the neural explants.

### Shh-P450 functions as a mitogen

In our neural explant assay, we noticed that Shh-P450 treated explants appeared to be larger than those treated with Shh-P150. To further investigate this, we performed a time course, tracking the change in size of individual explants, so that we could control for the variability introduced by dissections. Explants were incubated with 4 nM Shh-P150 (Fig. 3A), 4 nM Shh-P450 (Fig. 3B), 4 nM Shh-P150 with 1 μM Cyclopamine (Fig. 3C), 4 nM Shh-P450 with 1 μM Cyclopamine (Fig. 3D) or the respective HEK-derived exosome controls (Fig. S7). Explants were imaged using phase-contrast microscopy at the time of embedding in collagen (0 hr), followed by 24 hrs, and 48 hrs after incubation at 37^0^C. The area of each explant was determined and plotted as linked graphs (Fig. 3E-F) or normalized to their size at 0 hr (Fig. 3G). At 24 hrs and 48 hrs, the total area of explants treated with 4 nM Shh-P450 was significantly larger than the 4 nM Shh-P150 treated explants (Fig. 3G). This increase was inhibited by cyclopamine.

**Fig. 3:**
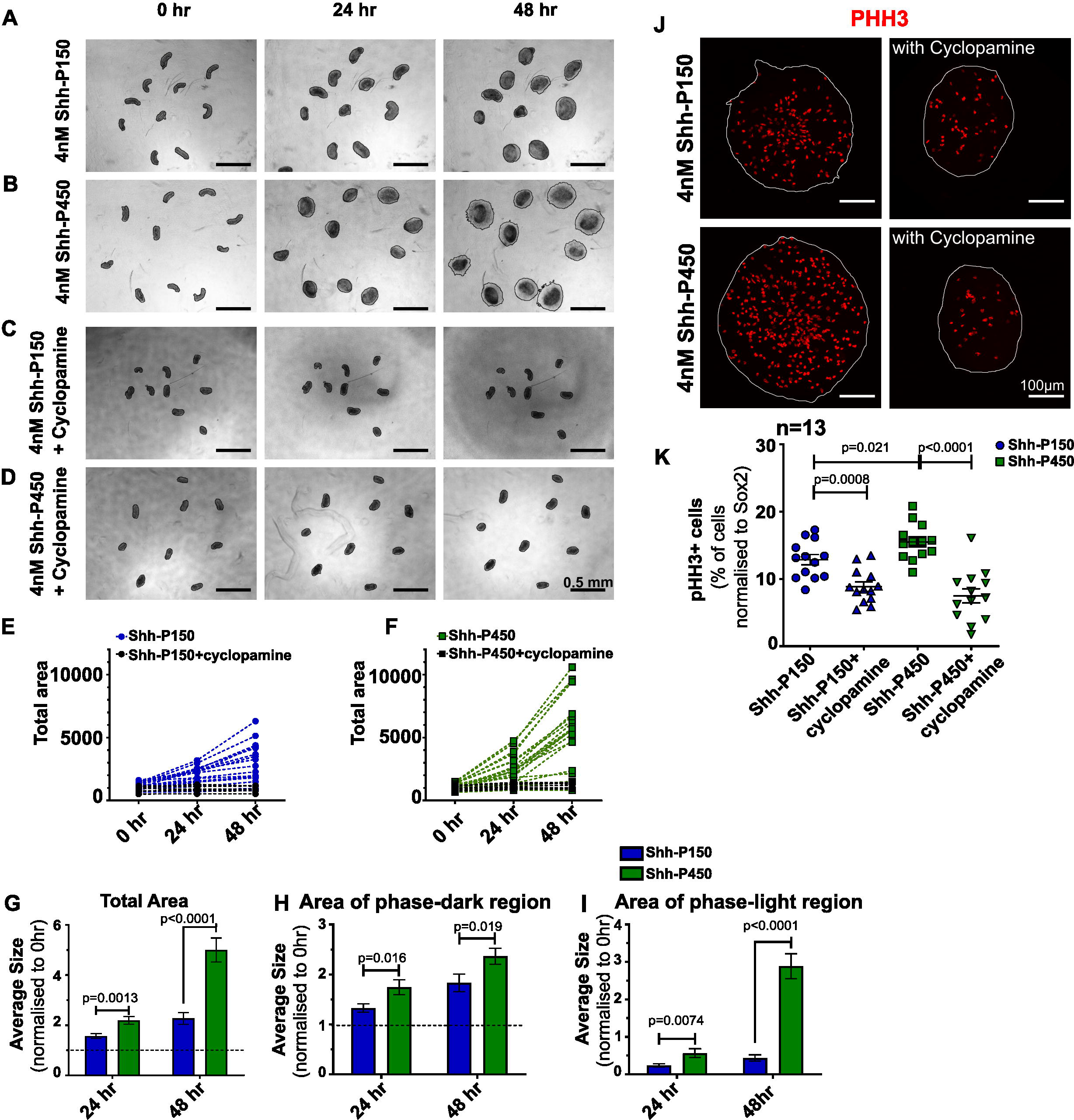
Shh-P450 promotes explant growth. (A-D) Representative phase contrast images of neural explants treated with 4 nM Shh-P150 (A), 4 nM Shh-P450 (B), 4 nM Shh-P150 with 1 μM cyclopamine (C) and 4 nM Shh-P450 with 1 μM cyclopamine (D). Images were taken at the time of embedding – 0 hr (left panel), after incubation at 37^0^C for 24 hrs (middle panel) and 48 hrs (right panel). Scale bar = 0.5 mm (E-F) Line graph representing change in the total size of each explant treated with 4 nM Shh-P150 (E, blue, n = 20), 4 nM Shh-P150 with 1 μM cyclopamine (E, black, n = 9), 4 nM Shh-P450 (F, green, n=20), and 4 nM Shh-P450 with 1 μM cyclopamine (F, black, n=9) across time – 0 hr, 24 hr and 48 hrs. n, number of explants. (G-I) Bar-graphs represent increase in the average size of the entire explant (G), inner phase-dark region (H), and outer phase-light region (I), normalized to the respective explant size at 0 hr, treated with 4 nM Shh-P150 (blue bars in G,H, and I), or 4 nM Shh-P450 (green bars, in G, H, and I), at 24 and 48 hrs. Dotted line (G and H) represent 0 hr. Statistical significance determined using Student’s T-test. Error bars represent mean ± SEM. (J) Representative images of explant stained with phospho-Histone H3 (pHH3) after treatment with 4 nM Shh-P150 or 4 nM Shh-P150 with 1 μM cyclopamine (top panel), and 4 nM Shh-P450 or 4 nM Shh-P450 with 1 μM cyclopamine (bottom panel) for 48 hrs. (K) Quantitative analysis of the percentage of pHH3 positive nuclei normalized to respective Sox2 positive nuclei, in the explants treated with 4 nM Shh-P150 (n=13), 4 nM Shh-P150 with 1 μM cyclopamine (n=13), 4 nM Shh-P450 (n=13), or 1 μM with 1 μM cyclopamine (n=13), for 48 hrs. Each dot represents data from one explant. Statistical significance determined using Student’s T-test. Error bars represent mean ± SEM. n, number of explants.

The Shh-P450 explants also showed a halo of phase-light cells (Compare Fig. 3A & B, 48 hrs) around the phase-dark core of cells. We hence evaluated the phase-light and phase-dark regions of these explants independently (Fig. 3H, I). Shh-P450 promotes the growth of the phase-light region more efficiently, which is also sensitive to cyclopamine (compare Fig. 3B & D). The phase-light region observed in Shh-P450 treated explants is not observed in HEK derived P150 or P450 treated explants (Fig. S7).

To further investigate if the larger explant area is a result of increase in cell division, we used phospho-Histone H3 (pHH3) to identify cells in late G2 and mitotic phase of cell cycle (Juan *et al*., 1998). Explants treated with 4 nM Shh-P450 for 48 hrs, show more pHH3 positive cells than 4 nM Shh-P150 explants (Fig. 3J-K). Cyclopamine reduced the size of the explants and the numbers of pHH3 positive cells in both Shh-P150 and Shh-P450 treated explants. This data suggests that Shh-P450 is a potential mitogen, supporting the proliferation of neural explants (Fig. 3J-K).

Overall our functional assays suggest that, the morphogenic and mitogenic abilities of Shh are segregated on Shh-P150 and Shh-P450 pools respectively.

### Regulation of Shh-P150 versus Shh-P450 secretion

Our data suggests that the two Shh exosomal pools have different functions. Identifying the regulators of these pools would provide insights into the physiological relevance of these activities. Our previous work had identified that proteins such as Flotillins, Integrins, Annexins were enriched in P150 over the P450 pool (Vyas *et al*., 2014). These proteins are generally sorted to membrane microdomains (De Gassart *et al*., 2003; Parchure, Vyas and Mayor, 2017). This may suggest that two pools could be derived from the same Multi-Vesicular Body (MVB), formed because of lipid segregation on MVB-limiting membrane. Lipids from Shh-P150, Shh-P450, HEK-P150, and HEK-P450 pools were extracted and subjected to LC-MS/MS analysis (Abhyankar *et al*., 2018; Pathak *et al*., 2018). We observed that when normalized to HEK-P150, the total lipid content of both Shh-P150 and Shh-P450 pools were similar (Fig. 4A & Table S1). Surprisingly, the presence of Shh on either P150 or P450 pools did increase the content of Ceramide, Cholesterol, Free Fatty Acid (FFA) and Sphingomyelin (SM). In contrast, the content of Phospholipids such as Phosphatidylcholine (PC), Phosphatidylethanolamine (PE) and Phosphatidic acid (PA) were reduced. However, when lipid content of Shh-P150 and Shh-P450 pools was compared with each other, there was no significant difference. This finding indicates that P150 and P450 do not result from differential lipid segregation on the MVB limiting membrane.

**Fig. 4:**
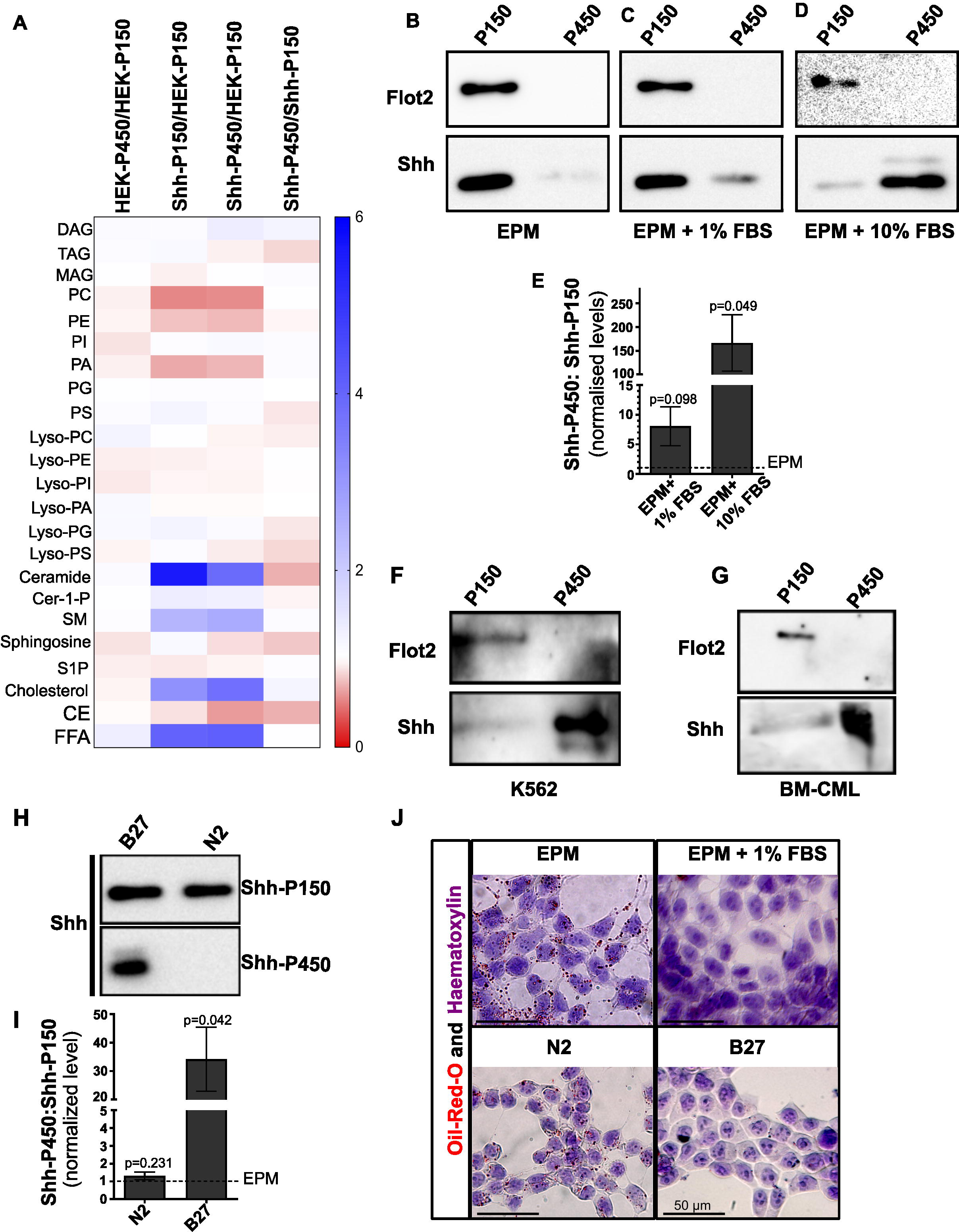
Regulators of Shh-P150 versus Shh-P450 pool. (A) Heat map represents relative levels of lipids, of Shh-P150, Shh-P450, HEK-P150 and HEK-P450 pools, normalized to HEK-P150 or as indicated. Red colour indicates downregulation and blue colour indicates upregulation in fold change. n= 6 biological replicates. (B-D) Representative western blots of Shh-P150 and Shh-P450 pools derived from Shh-HEK cells, cultured in EPM (B), EPM+1%FBS media (C) and EPM+10%FBS media (D), stained for Flot2 (top panel) and Shh (bottom panel). Equal total volume was loaded for P150 and P450 respectively. (E) Densitometric analysis to evaluate relative levels of Shh in B, C, and D. Ratio of Shh-P450 to Shh-P150 was derived for each condition after normalization for their respective loading volume and total protein concentration. Dotted line represents the normalized Shh-P450 to Shh-P150 ratio for EPM. n=3 biological replicates; Statistical significance determined using Student’s T-test. Error bars represent mean ± SEM. (F) Representative western blots of Shh-P150 and Shh-P450 exosomes derived from K562 cells transfected with Shh-cDNA and cultured in RPMI media supplemented with 10%FBS. The samples were probed for Flot2 (top panel) and Shh (bottom panel). (G) Representative western blots of P150 and P450 fractions derived from Human CML patient’s bone marrow derived plasma sample. Total three different samples were analysed. Equal protein was loaded per well and probed for Flot2 (top panel) and Shh (bottom panel). (H-I) Representative western blots for Shh on exosomes derived from HEK cells overexpressed with Shh, cultured in media supplemented with N2 or B27 defined supplements. (I) Bar-graph represents relative ratio of Shh-P450 to Shh-P150 pool derived from N2 or B27 supplemented media. Relative levels of Shh in each pool, was normalized to the loading volume and total protein. Dotted line represents the normalized Shh-P450 to Shh-P150 ratio for EPM. n=3 biological replicates; Statistical significance determined using Student’s T-test. Error bars represent mean ± SEM. (J) Representative images for Oil-Red-O (Red colour) and Haematoxylin (Blue colour) staining of cells cultured in different nutrient conditions-EPM (left; top panel), 1X DMEM with 1X N2 supplement (left; bottom panel), EPM with 1% FBS supplemented media (right; top panel) and 1X DMEM with 1X B27 supplemented media (right; bottom panel). n = 3 biological replicates.

Previous studies have suggested that serum acts as a regulator of Shh secretion (Panáková *et al*., 2005). Thus, we set out to understand whether the distinction of Shh-P150 or Shh-P450 secretion could be influenced by serum. For this, cells were cultured in exosome production medium (EPM) with or without serum. We find that by supplementing EPM with different amounts of serum, there was a proportional increase in Shh-P450 secretion with a corresponding decrease in Shh-P150 secretion (Fig. 4B-E). To probe if this was a specific feature of HEK293T cells, we checked if a chronic myeloid leukemia (CML) cell line, K562, would show a similar response. K562 cells transfected with Shh-cDNA, secrete Shh as P450 exosomes when cultured in 10% FBS supplemented media (Fig. 4F, K562). Shh is known to be secreted in CML patients bone-marrow (BM) aspirate on exovesicles (Anusha *et al*., 2021). Therefore, we analysed platelet-free plasma from CML patient’s bone-marrow aspirates. Here, again we find that Shh is present predominantly in P450 pool in bone-marrow aspirates (Fig. 4G, BM-CML), suggesting that this newly identified pool can be derived from different cell types and is also physiologically relevant.

To identify other culture conditions that could influence the secretion of Shh-P150 and Shh-P450 pools, we cultured Shh-HEK cells in EPM containing serum-free supplements, N2 or B27. We found that while N2 culture condition recapitulates the secretion profile of EPM, cells supplemented with B27 exhibit enhanced P450 secretion (Fig. 4H-I).

The overlapping secretion profiles triggered by EPM and N2 versus Serum and B27, suggested that these culture conditions might be inducing similar cellular states. Lipid droplet (LD) accumulation has been reported to signify nutritionally starved cells (Schroeder *et al*., 2015). To explore if our culture conditions correlated with starvation, we used Oil Red O, a marker of LDs, together with the nuclear dye Haematoxylin. We find that LDs were present in Shh-HEK cells (Fig. 4J, left panel) cultured in EPM or N2 supplemented condition representing a starved state. In contrast, LD content was reduced or nearly absent in serum or B27 supplemented media, representing a fed state (Fig. 4J, right panel).

### Rab7 is required for Shh-P150 secretion

Our results suggest a correlation between LD accumulation and Shh-P150 secretion. The small GTPase Rab7 has previously been shown to be associated with LD accumulation (Schroeder *et al*., 2015). We thus asked whether Rab7 might be responsible for Shh-P150 secretion. We used the proteomics analysis of P150 and P450 pools derived from HEK293T and Shh-HEK cells that was performed in our previous study (Vyas *et al*., 2014) to gain insights in to the biogenesis of these two pools. By reanalyzing the vesicular transport proteins in this data (Fig. 5A), we find that both P150 and P450 pools shared common early endosomal regulators, such as Rab1a, Rab6a, Rab8a, Rab8b, Rab10, Rab14, and Rab35 (Fig. 5A, Table S2). However, Shh-P150 showed the specific association of late endosomal and MVB biogenesis markers such as Rab7a, Rab13, Rab21, Ral-A, CHMP2B, CHMP4B, Syntaxin-7, etc. (Fig. 5A, Table S2). We measured the relative levels of the early endosomal markers, Rab5 or EEA1 and late endosomal marker, Rab7 or the ESCRT protein TSG101, in cell lysates obtained from Shh-HEK cells cultured in EPM or EPM + 1% serum. Rab5, EEA1 and TSG101 levels showed no significant differences in the two culture conditions (Fig. 5B). Consistent with earlier reports (Schroeder *et al*., 2015), Rab7 expression was higher in Shh-HEK cells cultured in only EPM (Fig. 5B).

**Fig. 5:**
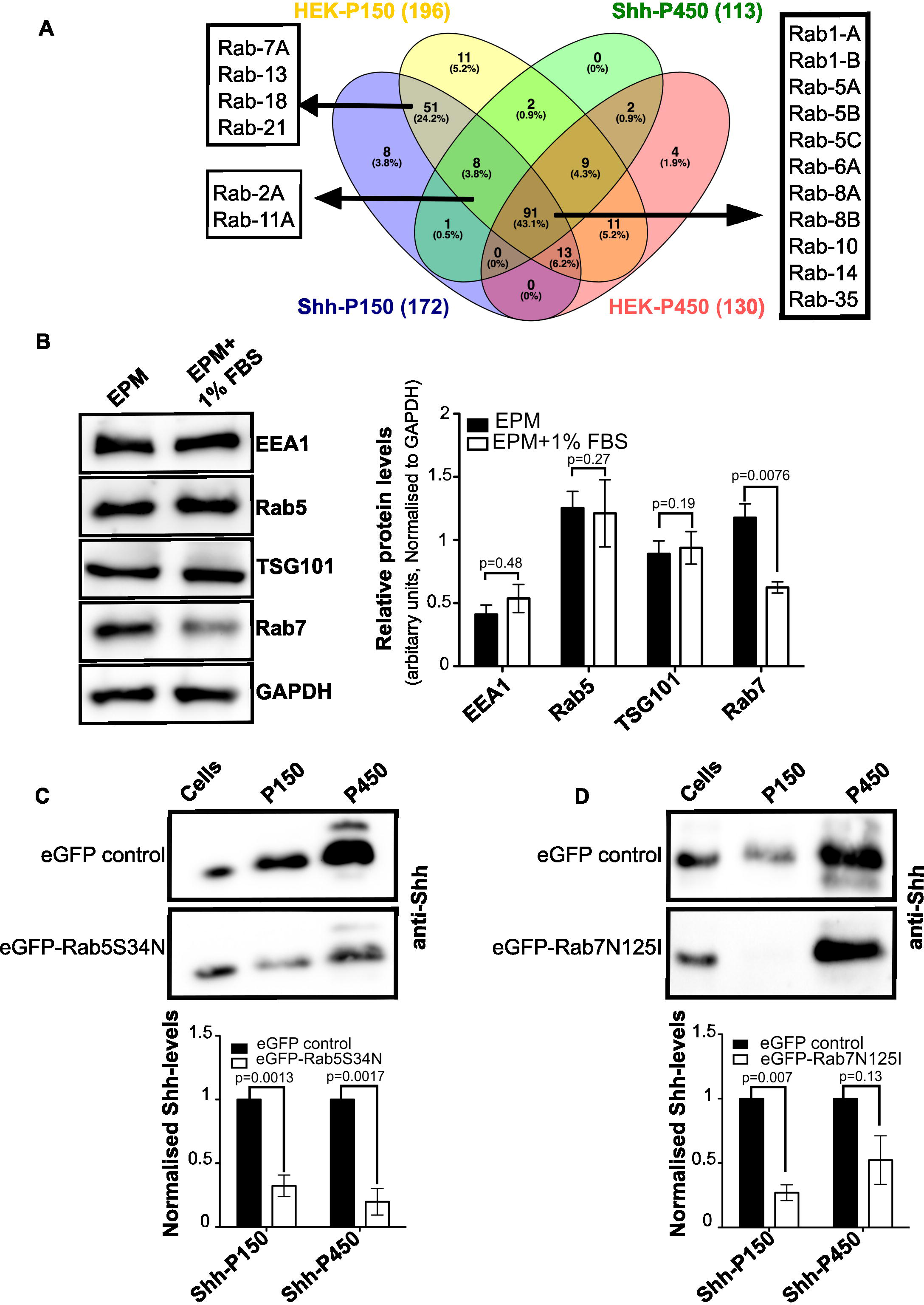
Rab7 regulates secretion of Shh-P150 pool. (A) Venn diagram representing vesicular transporter proteins identified using proteome discoverer^TM^ software 1.4 present in the different exosomal pools, HEK-P150 (yellow), Shh-P150 (blue), HEK-P450 (red) and Shh-P450 (green). The numbers in brackets represent the total number of vesicular transporters respectively. (B) Representative western blots of EEA1, Rab5, TSG101, Rab7, and GAPDH on cells cultured in EPM or EPM with 1% FBS media. Densitometric analysis of the blots presented as bar graph normalized to respective GAPDH levels. n=3 biological replicates; p-value determined by Student’s T-test. Error bars represent mean ± SEM. (C-D) Representative western blots showing levels of Shh in P150 or P450 pools, derived from Shh-HEK cells transfected with eGFP vector (control, top panel, C and D) dominant negative eGFPRab5S34N (bottom panel, C), or dominant negative eGFPRab7N125I (bottom panel, D). Blots were probed with anti-Shh antibody. The graphs represent densitometric analysis of the western blots, presented as fold change normalized to the respective eGFP controls. n=3 biological replicates; Statistical significance determined using Student’s T-test. Error bars represent mean ± SEM.

The above restriction of expression suggested that while both Shh pools require the early endosomal protein Rab5, Rab7 was specifically required for Shh-P150 maturation/secretion. To evaluate this possibility, we perturbed Rab5 or Rab7 using overexpression of GFP - tagged dominant negative forms of these proteins. eGFP overexpression was used as control for these assays. Shh-HEK cells expressing a dominant negative form of Rab5 (eGFP-Rab5S34N) significantly reduced secretion of both the vesicular pools (Fig. 5C). In contrast, Shh-HEK cells transfected with a dominant negative Rab7 construct (eGFP-Rab7N125I) show much reduced secretion of Shh-P150 exosomes, while Shh-P450 secretion was not significantly affected (Fig. 5D). This suggests that while Rab5 can regulate the biogenesis of both Shh-P150 and Shh-P450 pools, Rab7 specifically regulates Shh-P150 production.

### Rab7 is essential for Shh-mediated ventral neural tube patterning

During neural tube patterning Shh is secreted by the notochord to specify ventral fate in the neural tube. Our explant assays show that ventral neural tube patterning is mediated through Shh-P150 exosomes. Furthermore, we find that Rab7 regulates Shh-P150 biogenesis in cell lines (Fig. 5D). We thus asked if we could use these findings to understand the necessity of Shh-P150 function in neural tube patterning. We first used immunohistochemistry to evaluate endogenous Rab7 levels in notochord and developing neural tube using transverse sections of chicken HH-17 stage embryos. We detected robust Rab7 expression in the notochord when compared to adjacent neural tube tissue (Fig. 6A). This finding is consistent with previous reports on active late endosomal biogenesis in the notochord (Ellis, Bagwell and Bagnat, 2013; Kreis, Wielath and Vick, 2021).

**Fig. 6:**
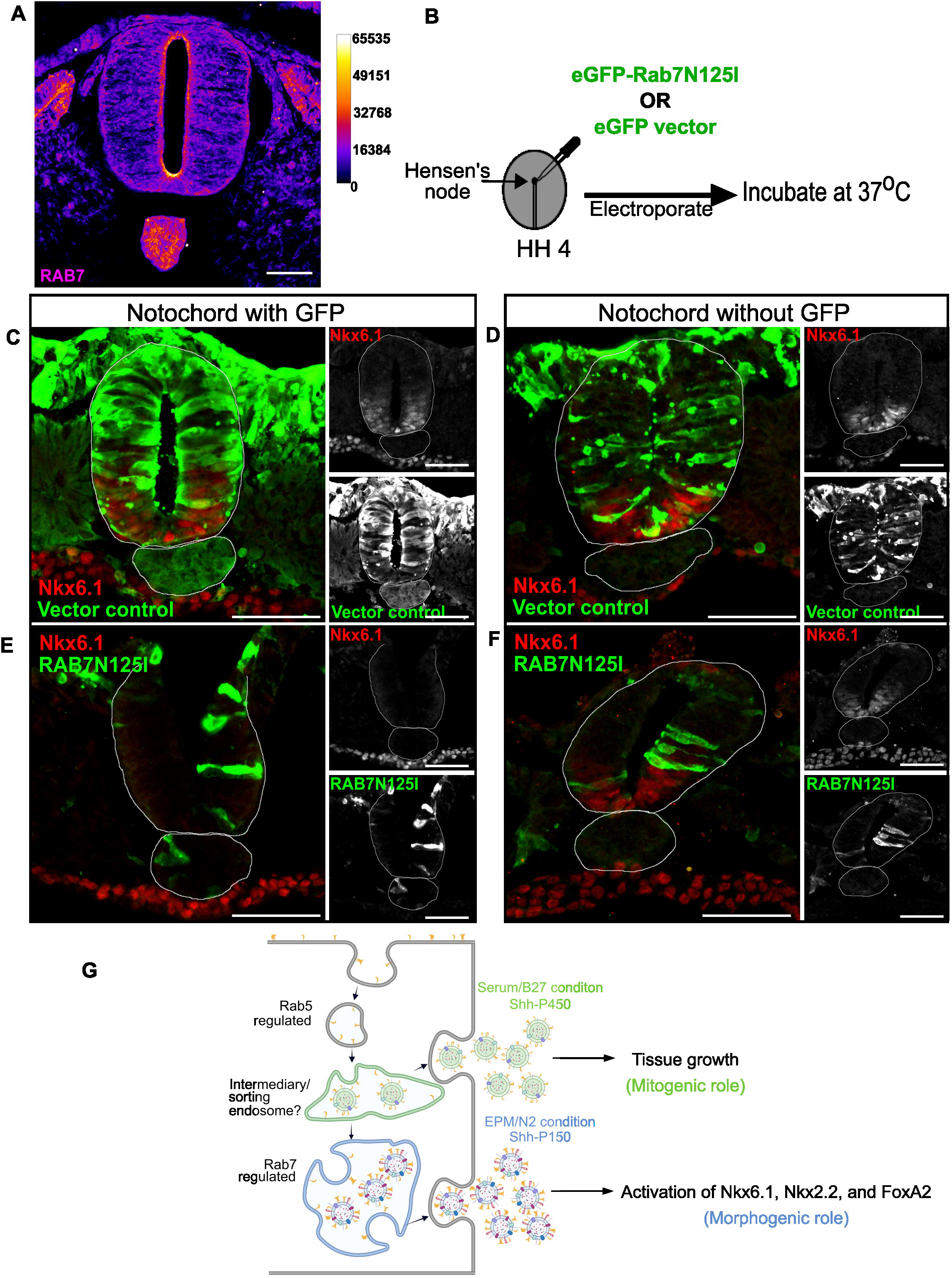
Rab7 is essential for Shh-mediated ventral neural patterning. (A) Representative image of endogenous Rab7 staining on transverse sections of HH-17 stage chicken embryo. The image is a FIRE LUT representation that distinguishes between higher and lower intensity pixels. Scale bar = 50 μm. The LUT bar represents the intensity values. (B) Schematic representation of electroporation procedure performed at the Hensen’s node of HH-4 stage chicken embryo. (C-F) Representative immunostaining of transverse sections of embryos electroporated with eGFP vector (C, D, n=3) or eGFP-Rab7N125I (E, F, n=5). For eGFP-vector electroporated embryos, Nkx6.1 expression (red in C or D) is observed in absence or presence of notochordal-GFP (Green in C or D). For eGFP-Rab7 electroporated embryos, neural tube Nkx6.1 expression (red in E or F) can be observed only in absence of notochordal-GFP (green in E or F). Insets on the right of each image represent individual fluorescence channels, respectively. n, number of embryos. Scale bar = 100 μm. (G) Schematic depicting mechanism of Shh exosome biogenesis and their function. Rab5 regulates biogenesis of Shh-P150 and Shh-P450 whereas Rab7 regulates biogenesis of Shh-P150 pool. Shh-P450 pool may be derived from intermediate or sorting endosomes. Image created with BioRender.

To probe the role of Rab7 in ventral neural tube patterning, we over expressed a GFP-tagged dominant negative Rab7 cDNA in the notochord. For this, either a control eGFP plasmid or eGFP-Rab7N125I plasmid was electroporated into Hensen’s node in HH4 stage embryos. These embryos were then incubated for 24 hrs at 37°C (Fig. 6B). Embryos were then sectioned and both GFP and the ventral neural tube marker, Nkx6.1 were probed. We observed mosaic expression of eGFP-vector (n=3; Fig. 6C-D) or eGFP-Rab7N125I transgene (n=5; Fig. 6E-F) in the notochord. This feature allowed us to evaluate sections with and without notochordal GFP expression from the same embryos. Regions where notochordal cells were positive for eGFP-Rab7N125I, showed a down-regulation of Nkx6.1 (Fig. 6E). Endodermal Nkx6.1 expression, which is independent of the notochordal Shh (Hebrok, Kim and Melton, 1998), remained unaffected in these regions (Fig. 6C-F). Interestingly, the regions where eGFP-Rab7N125I was not expressed in the notochord, Nkx6.1 expression was present in the neural tube (Fig. 6F). These data suggest that Rab7 perturbation in notochord locally influences neural tube Nkx6.1 expression. Overall, our data here suggests that active notochordal Rab7 is necessary for ventral neural tube patterning, likely through the secretion of Shh-P150.

## Discussion

In this study we show that Shh carried on two pools of exosomes and isolated by differential centrifugation, can have distinct signaling characteristics in the neural tube. The patterning activity of Shh is segregated on the classical exosomal pool, Shh-P150; while a lighter exosomal pool, Shh-P450, is responsible for the mitogenic activity of Shh. Importantly, we suggest that secretion of one form over the other can be spatially restricted, with Rab7 function important for the secretion of Shh-P150 from the notochord.

Shh-P150 can induce protein expression of all ventral markers of the spinal cord. Whereas, Shh-P450 drives proliferation and can also induce transcript of the ventral progenitor gene Nkx6.1. In both cases, these activities are sensitive to the inhibition of Smoothened. This does suggest that the engagement of exosomal Shh with Patched, to relieve the inhibition of Smoothened is unaffected. The specificity of Shh response is likely to be driven by the specific components present on or in the exosomes.

### Cofactors carried on exosomes

Our previous work had identified numerous P150 or P450 specific proteins and miRNAs that may be important in encoding the specificity of the Shh response (Vyas *et al*., 2014).

In absence of Shh, its receptor Patched inhibits a GPCR, Smoothened (Ingham *et al*., 2000). Upon Shh binding to Patched, this inhibition is relieved. It then activates the nuclear effectors of Shh signaling, the Gli proteins, by blocking their repressor, protein kinase A (PKA) (Ayers and Thérond, 2010). Two possible mechanisms are thought to link Smoothened with PKA. The first is canonical, with trimeric G proteins activity modulating the activity of cyclic AMP (cAMP), which impacts the activity of PKA (Pal and Mukhopadhyay, 2015). In recent years, an alternate, direct link with PKA activity has also been identified (Arveseth *et al*., 2021). One possible mechanism that could determine the specificity of the Shh response may be the switch between these two modes of PKA regulation. Indeed Shh-P450 vesicles exclude G protein subunit α i2 (GNAI2), which is found in HEK-P150, Shh-P150 and HEK-P450 exovesicles (Table S2). These could act downstream of either Smoothened, or other GPCRs that have been shown to influence Shh signaling, such as Gpr161 (Pusapati *et al*., 2018), Gpr175 (Singh, Wen and Scales, 2015) or Gpr17 (Yatsuzuka *et al*., 2019). It is possible that local modulation of the Smoothened/GPCR response through the delivery of specific Gα protein subunits, or by other means, is sufficient to change the specificity of signaling. One outcome of this switch could be the cohort of Gli genes that are active. This idea is supported by our observations that the treatment of explants with either Shh-P150 or Shh-P450 shows upregulation of Gli1, however only Shh-P450 shows upregulation of Gli2 expression, perhaps indicative of differences in PKA activation downstream of Shh-receptor engagement.

While our analysis did not detect specific changes in lipid composition between Shh-P150 and Shh-P450 exosomes, we did detect different membrane proteins between the two. Again, it is possible that these co-receptors allow the assembly of distinct Shh receptorsome complexes, with one of the Shh co-receptors; HH interacting protein (HHIP) (Chuang and McMahon, 1999), Cell adhesion molecule-related/down-regulated by oncogenes (Cdon), Brother of Cdon (Boc) or Growth-arrest specific protein-1 (Gas1) (Allen *et al*., 2011). We detect Integrin-β1 on Shh-P150 which may aid in targeting Shh-P150 to specific lipid microdomains with distinct receptor/co-receptor complexes, and by extension distinct downstream transducers (Vyas *et al*., 2014). Shh-P450 shows specific co-expression of the Low Density Lipoprotein (LDL) receptor family protein, LRP2 (Table S2). LRP2 also acts as a Shh co-receptor, promoting uptake of the Shh-Patched complex (Christ *et al*., 2012). Again it is possible that altering the way in which the Smoothened inhibition is lifted enables a toggle in its downstream activation pathway.

The block in the translation of Nkx6.1 suggests that Shh-P450 may also carry cargoes important in post-transcriptional regulation of protein production. Our previous studies had also identified a number of specific microRNAs that are found in these two exosomal pools. Shh-P450 in particular carries specific miRNAs (Vyas *et al*., 2014). Homology searches show that they have targets in ventral neural tube markers such as Nkx6.1 and Nkx2.2 (Fig. S8). As proposed (Vyas and Dhawan, 2017), co-sorting of such partners in the exosomal lumen can add a further level of control in the response that Shh-P150 and Shh-P450 evoke on neural tissue.

Our findings here suggest that packaging of Shh with different co-factors/partners might be involved in modulating different signaling abilities of Shh. Such mechanisms can ensure that despite limited repertoire of signaling proteins, the permutation and combination of other critical partners can influence the cellular signaling networks at different nodes and there by generate distinct outcomes to pattern the developing tissue.

### Spatial regulation of exosomal biogenesis

The role of Shh in ventral patterning of the neural tube is now well established. Here Shh is secreted from the notochord acting on adjacent neural tissue to mediate these effects (Roelink *et al*., 1995; Ribes and Briscoe, 2009; Sagner and Briscoe, 2019). We found that Rab7 regulated the Shh-P150 exosomal pool. This is consistent with the high Rab7 levels in the notochord and its role in ventral patterning within the neural tube. It is unclear what is the source of Shh-P450 signals, however proliferation in much of the neural tube occurs in the ventricular, luminal zone (Cayuso *et al*., 2006) and it is likely that P450 vesicles act in this region. It is possible that relative difference in Rab7 levels influences endosomal maturation rate thereby regulating biogenesis of Shh-P150 versus Shh-P450.

The mitogenic role for Shh-P450 is consistent with our previous study where we found that, in CML patient plasma, exovesicular Shh confers resistance to therapy (Anusha *et al*., 2021). We show that the dominant pool of Shh secreted in CML patient samples is the Shh-P450 pool, and it is possible that the role of Shh-P450 may have an underappreciated role beyond the developmental context we have investigated.

Overall, our study suggests that P150 and P450 are exosomes derived by differential endosomal maturation or sorting mechanism (Fig. 6H). Segregation of Shh with different partners thus influences its signaling abilities (Fig. 6H). Apart from tissue patterning, Shh is also implicated in different metabolic disorders (Garg *et al*., 2022) and cancers (Barakat, Humke and Scott, 2010). More concerted efforts are required to resolve the critical partners of Shh in different conditions to interpret the complex regulation mediated by Shh, providing ways in which the distinct activities of Shh can be modulated in development and disease.

## Acknowledgements

N.V. and A.W. would like to thank Dr. Anna Kicheva for discussions regarding the explant assay. N.V. thanks Dr. James Briscoe for introducing her to the explant assay technique. We would like to thank Neelay Mehendale and Saddam Shaikh for technical support and assistance for lipid profiling. NCBS-Central Imaging and Flow Facility (CIFF) for Confocal microscopy. CPDOTI for providing fertilized chicken eggs.

## Author contributions

A.W: performed research, involved in designing experiments, analyzed data, data interpretation, generated data figures, and manuscript writing.

S.P: Electroporation, cryosectioning and analysis.

S. S K: Sample processing for LC-MS and lipid profile analysis, data acquisition and interpretations.

S.D: provided clinical samples, clinical data accrual, and regulatory permission.

R. K. L: Data interpretation and manuscript writing.

N.V: Study design, fund raising, data interpretation, and manuscript writing.

## Materials and Methods

### Isolation of exosomes

HEK293T cells with or without Shh cDNA transfection were used for exosome isolation. Cells were cultured for 2 days in DMEM media containing 10% FBS, to achieve ∼70% confluency. These cells were then washed with Phosphate Buffered Saline (PBS) and supplemented with Exosome Production Media (EPM) (Vyas *et al*., 2014) or EPM supplemented with different percentages of FBS and further incubated for 48 hrs. After 48 hrs, the conditioned media was collected and cleared for cellular debris by differential centrifugation. The supernatant was subjected to ultracentrifugation to isolate exosomes using tabletop ultracentrifuge (Optima^TM^ MAX-XP) in TLA100.3 rotor. The exosomal pellet obtained after 150,000g ultracentrifugation spin for 1.5 hrs at 4^0^C was designated as P150. The supernatant obtained was further subjected to ultracentrifugation spin at 450,000g spin for 1.5 hrs at 4^0^C and the pellet obtained was designated as P450. P150 and P450 pellets were dissolved in equal volume of 1X PBS for further analysis.

### Neural plate explant culture

Fertilized chicken eggs were incubated in a humidified incubator at 37^0^C for 36-38 hrs until they reached stage HH10 (10 somite stage). The neural plate caudal to the somites was dissected and placed in L15 medium (Gibco). The dissected neural plate tissue was treated with dispase (10 mg/ml in L15) on ice for 15 min. The neural plate tissue was obtained by removing the subjacent mesoderm, notochord and other associated tissue. The neural plate was cut to appropriate sizes and stored in L15 on ice, until they were embedded. Collagen substrate was prepared with 400 μl of 3 mg/ml rat-tail collagen (Gibco) in 5 μl 1M HEPES (pH 7) supplemented with 50 μl 10X DMEM and 30 μl 7.5% Sodium bicarbonate to adjust the pH. Collagen droplets were used as a substrate to embed the explants in 4 well dishes. The explants were placed in the collagen droplets and incubated at 37^0^C in 5% CO_2_ incubator until the droplet solidified and then different conditions were added onto the explants in Advanced DMEM-F12 media (Gibco) supplemented with 1X N2 (serum free supplement) (Gibco), 1X PenStrep and 1X Glutamax and incubated for 24 or 48 hrs at 37^0^C.

### Cryosectioning

Embryos of required stage were fixed in 4% PFA overnight at 4^0^C. Upon fixation, embryos were thoroughly washed with PBS and equilibrated with 15% sucrose overnight at 4^0^C. The embryos were then embedded in 7.5% gelatin and frozen at −20^0^C. 30 μm serial sections of the thoracic region of the embryo were obtained using a cryostat (Thermo Scientific HM525NX). Antibody staining was performed on these cryosections.

### Immuno-staining

The explants were fixed in 4% PFA, then permeabilised with 0.1% Triton X-100 for 15 min at RT and then blocked using 2 mg/ml BSA. Primary antibodies were diluted in block solution and incubated overnight at 4^0^C (Refer Table S3 for antibody details). Upon incubation the primary antibody was washed off with PBS + 0.1% TritonX-100 (PBST) thrice. Secondary antibody was diluted in block and added onto the explants and incubated for 1 hr at room temperature (RT). Secondary antibody was washed off with PBST and the explants were mounted onto glass slides and images acquired using confocal microscope (Olympus FV3000).

For cryosections – The sections were permeabilized with 0.5% Triton X-100 for 10 min at RT and blocked with 2 mg/ml BSA blocking solution. Primary antibodies in block were prepared and added onto the slides (Antibody details in Table S3). The slides were kept in a humidified chamber at 4^0^C overnight. This was followed by washes and secondary antibody (Table S3) incubation for 1.5 hrs at RT. The secondary was washed off with PBST and the slides were mounted and imaged using fluorescence microscope (Leica DM 5000B) or confocal microscope (Olympus FV3000).

### Image Analysis

Images acquired were analyzed on Imaris 9.3 software (Oxford Instruments). The explants were imaged with same imaging settings. The spots function was used to determine the number of Sox2+ nuclei or ventral neural progenitor (FoxA2, Nkx2.2 and Nkx6.1) nuclei. Percentage of the ventral progenitor markers was determined with respect to the Sox2 nuclei for each explant.

### RNA isolation and cDNA synthesis

The explants/cells were suspended in TRIzol reagent (Invitrogen) and RNA was extracted. The samples were subject to chloroform treatment (20%) and incubated at room temperature for 5 mins. The samples were then centrifuged at 13,000 rpm at 4^0^C for 15 mins. The aqueous phase was separated and equal volume of isopropanol was added. Samples were incubated at room temperature for 10 mins and centrifuged at 13,000 rpm at 4^0^C for 15 mins. The pellets obtained were washed with 75% ethanol, air-dried, and resuspended in nuclease-free water. 1-2 μg total RNA was used for cDNA synthesis. cDNA was synthesized using the Applied Biosystems High Capacity cDNA Reverse Transcription Kit according to the manufacturer’s instructions.

### Quantitative PCR (qPCR)

The cDNA synthesized was followed with Real Time-quantitative PCR using Applied Biosystems SYBR Green PCR Master Mix. qPCR was performed using QuantStudio^TM^ 6 Flex Machine (Thermo Fischer Scientific). Fold change was calculated using the 2(–ΔΔCt) method compared to the loading control GAPDH (primer sequences listed in Table S4)

### Protocol for solubilizing lipids - Lipid extraction

Exosomes were isolated and suspended in 1 ml molecular grade PBS (Phosphate Buffer Saline), vortexed and transferred to a glass tube. To this, 3 ml of 2:1 chloroform: methanol containing different internal standards (Avanti Polar lipids) reported earlier (Abhyankar *et al*., 2018; Pathak *et al*., 2018) (Table S5) was added, and this mixture was vigourously vortexed, and the organic layer was separated as discussed earlier. The phospholipids were enriched by acidification of the remaining aqueous layer by addition of 1% (v/v) formic acid, and re-extracted using 2 ml of chloroform as reported earlier (Abhyankar *et al*., 2018; Pathak *et al*., 2018). All LC-MS analysis was done on a Sciex X500R fitted with an Exion UHPLC system using MRM-HR scanning method reported earlier (Abhyankar *et al*., 2018; Pathak *et al*., 2018). All the endogenous lipid species were quantified by measuring the area under the curve in comparison to the respective internal standards, and then normalized to the total protein content of the individual preparations.

### SDS-Polyacrylamide Gel Electrophoresis (SDS-PAGE) and Western Blotting

Cells or exosomes were dispensed in lysis buffer consisting of 300 mM NaCl, 50 mM Tris pH 7.4 and 0.5% Triton X-100. The lysed samples were used for protein estimation with Bradford reagent (Bio-Rad). The samples were boiled for 5 minutes at 95^0^C in 5X SDS loading dye and loaded on to 12% SDS-PAGE gels. Electrophoresis was performed at 120V (Bio-Rad). Once the proteins resolved, they were transferred onto nitrocellulose membrane (Pall corporation) overnight at 4^0^C at 20V. After transfer, the membrane was blocked for 1 hr at RT with 5% blotto prepared in Tris Buffer Saline with 0.1% Tween20 (TBST) followed by primary antibody incubation overnight at 4^0^C. The blot was washed with TBST and HRP-conjugate secondary antibody (Table S3) was incubated for 1.5 hrs at RT. After this the blot was imaged in ImageQuant^TM^ LAS 4000.

### Bone Marrow sample processing for exosome isolation

Leftover samples of Bone-marrow aspirates collected in EDTA-vacutainers for our previous study (Anusha *et al*., 2021), (IRB. No. NHH/AEC-CL-2017-172) were used here. Ethics waiver was taken from the institute review board (letter dated 5^th^ June 2023) to use these leftover derivatives for this study. Cells were removed from the bone-marrow aspirates by centrifugation at 1000 rpm, for 5 min. The resultant plasma fractions were collected in a separate tube and centrifuged at 2500 rpm for 5 min and then stored at −80^0^C. 50-100 μl of the stored samples were diluted in 2.5 ml of PBS and subjected to differential centrifugation at 5000 rpm for 15 min followed by another centrifugation of the supernatant at 13000 rpm for 30 min, to clear debris. The samples were then subjected to sequential ultracentrifugation at 150,000g and 450,000g (Optima^TM^ MAX-XP; TLA100.3 rotor). The resultant pellets were designated as P150 and P450 and were subjected to western blotting analysis.

### Oil-Red-O staining

Cells were cultured in different nutrient conditions. Upon 70% confluency the cells were washed with PBS and were fixed in 4% PFA. Following fixation, the cells were rinsed in distilled water treated with 60% isopropanol for 2 mins. Cell were then stained with Oil-red-O (Sigma-Aldrich) for 10 mins, followed by washing with 60% isopropanol, distilled water, and Haematoxylin (Sigma-Aldrich) staining for 2 minutes. The cells were rinsed twice until remnant Haematoxylin was washed off, mounted, and imaged under bright field on microscope (Olympus BX-51).

### Electroporation

Freshly fertilized hens’ eggs were incubated in a humidified chamber at 37^0^C for 18 hrs. Early embryos were staged using the Hamburger Hamilton stage series (HH; Hamburger and Hamilton, 1951). Rab7 knockdown was performed by electroporating a Rab7 dominant-negative construct (eGFP-Rab7N125I) while eGFP-C1 vector plasmid was used as control. HH4 chicken embryos were collected and placed in a Petridish plate electrode (CUY701P2E) containing Ringer’s solution. Plasmids were injected (plasmid mix for microinjection in Table S6) at the Hensen’s node of the embryos using a Microloader pipette tip (Eppendorf). Electroporation was performed using CUY21EDIT (Nepa Gene) by giving 5 pulses of 7V/0.01A, 100ms each. Electroporated embryos were cultured *ex ovo* on albumin-agar plates in a humidified CO_2_ incubator at 37^0^C for 24 hours. These cultured embryos were then fixed in 4% PFA overnight at 4^0^C for cryosectioning followed by immunohistochemistry (Antibody details in Table S3).

### Statistical Analysis

Data are represented as mean ± SEM. All statistical analyses were performed using GraphPad Prism 8.0.

